# Incorporating knowledge of plates in batch normalization improves generalization of deep learning for microscopy images

**DOI:** 10.1101/2022.10.14.512286

**Authors:** Alexander Lin, Alex X. Lu

## Abstract

Data collected by high-throughput microscopy experiments are affected by batch effects, stemming from slight technical differences between experimental batches. Batch effects significantly impede machine learning efforts, as models learn spurious technical variation that do not generalize. We introduce *batch effects normalization* (BEN), a simple method for correcting batch effects that can be applied to any neural network with batch normalization (BN) layers. BEN aligns the concept of a “batch” in biological experiments with that of a “batch” in deep learning. During each training step, data points forming the deep learning batch are always sampled from the same experimental batch. This small tweak turns the batch normalization layers into an estimate of the shared batch effects between images, allowing for these technical effects to be standardized out during training and inference. We demonstrate that BEN results in dramatic performance boosts in both supervised and unsupervised learning, leading to state-of-the-art performance on the RxRx1-Wilds benchmark.^1^

## 1 Introduction

In recent years, there has been great interest in using deep learning to analyze images collected using microscopy [Moen et al., 2019]. One area is morphological profiling, which seeks to identify changes in the morphology of cells in response to chemical or genetic perturbations. These morphological changes serve as a general read-out of phenotype, enabling downstream applications like drug development or functional genomics [Caicedo et al., 2016]. By pairing these principles with high-throughput microscopes [Usaj et al., 2016], biologists collect thousands of images of cells in parallel, each with a different treatment, enabling unbiased systematic discovery [Bray et al., 2016]. However, analyzing the large volume of images generated by these experiments requires computational strategies, leading to great interest in deep learning methods for morphological profiling [Pratapa et al., 2021, Hallou et al., 2021].

However, one central challenge in using deep learning (and computer vision in general) to analyze microscopy images is *batch effects* [Leek et al., 2010, Goh et al., 2017]. Technical variations in experimental condition (e.g. day-to-day differences in temperature or humidity, or differences in microscope settings) can lead to systematic differences in images that were collected in the same experiment [Caicedo et al., 2017]. While subtle to the eye, batch effects may be correlated with desired biological signal. As a result, neural networks can overfit to them as a surrogate for learning biological signal, leading to failures in generalization and incorrect conclusions in data analysis [Lu et al., 2019a, Ando et al., 2017].

In this work, we present *batch effects normalization* (BEN) – a simple method for correcting batch effects that can be applied to any deep neural network with batch normalization (BN) layers [Ioffe and Szegedy, 2015]. BEN relies on a simple tweak in training and inference: instead of drawing an input batch to a neural network randomly, we enforce that all samples come from the same batch of biological experiments (hereby referred to as an “experimental batch” to disambiguate from a “batch” in deep learning). When combined with BN layers, this sampling strategy allows the model to learn features that have been standardized to the rest of the experimental batch, removing batch effects (Figure 1).

**Figure 1:**
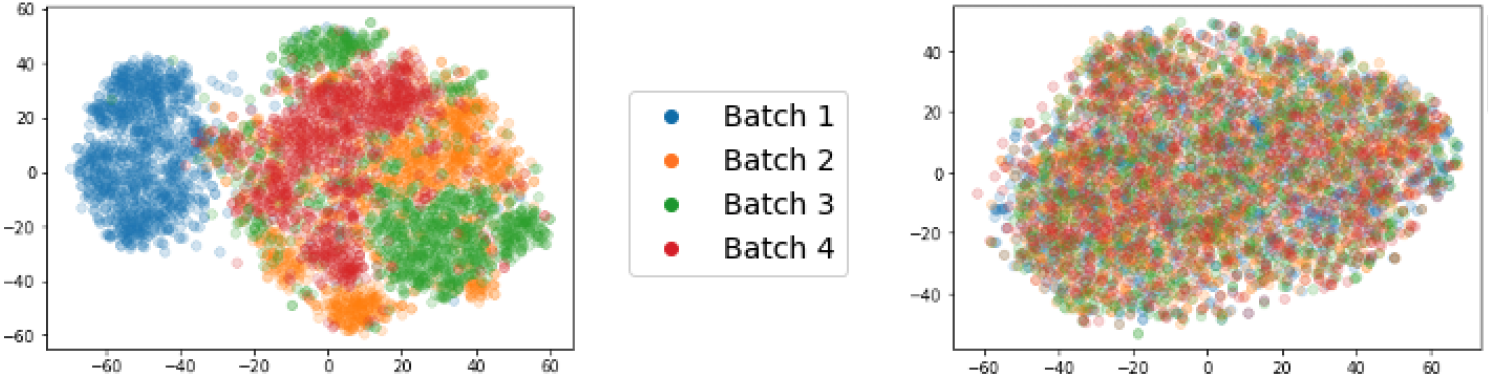
(Left) The representation space of microscopy images learned by a deep convolutional neural network with standard BN layers, projected to 2 dimensions using t-SNE. (Right) The representation space learned by the same network when BEN is applied. Further details can be found in Appendix B.1.2.

We show that on the RxRx1-Wilds dataset [Taylor et al., 2019, Koh et al., 2021], a challenging dataset used to study batch effects in microscopy images, BEN results in performance gains regardless of machine learning strategy, generically improving the performance of supervised learning, self-supervised learning, and transfer learning at both the single cell crop and whole image levels. Overall, BEN requires minimal changes to pre-existing deep learning pipelines, but results in significant mitigation of batch effects:

- In supervised learning, applying BEN to a standard ResNet classifier results in state-of-the-art performance, outperforming previous methods relying on highly engineered domain adaptation strategies (Section 4.1).
- In unsupervised learning, BEN nearly doubles the performance of an off-the-shelf self-supervised method previously shown to be effective for morphological profiling [Perakis et al., 2021]. We further propose a new self-supervised algorithm and show that BEN nearly doubles its performance as well, suggesting that BEN will generalize to novel methods in the future (Section 4.2).
- Transfer learning strategies are effective for morphological profiling, but to-date, models trained on microscopy images [Hua et al., 2021] have shown worse transfer performance than models trained on natural images [Ando et al., 2017]. BEN closes this gap, resulting in transfer learning performance on-par with the top natural image models for the BBBC021 dataset [Caie et al., 2010] (Section 4.4).

## 2 Method

In this section, we explain *batch effects normalization* (BEN), our method that corrects for experimental batch effects within any neural network architecture containing batch normalization (BN) layers. Modern biological experiments are often collected in multiple experimental batches. For example, in high-throughput microscopy, cells are cultivated in plates containing many wells, each hosting an individual experiment; images are taken of each well, but we consider all of these images to be of the same experimental batch because they come from the same plate.

To implement BEN, only three changes are required in the training and inference of a deep learning model:

1. *Batch-wise Training:* During training, each training batch exclusively contains samples drawn from a single experimental batch (as opposed to randomly from the entire dataset).
2. *Batch-wise Inference:* During inference, predictions are obtained by feeding data from an entire experimental batch to the model at once (as opposed to feeding each data point one-by-one).
3. *Unfrozen Batch Normalization:* During inference, the BN layers should not be frozen (i.e. instead of using historical statistics from the training dataset, the BN statistics should be re-calculated based on the inference batch.)

Classically, BN layers stabilize the training of neural networks by normalizing each feature by batch statistics (i.e. means and variances) [Ioffe and Szegedy, 2015]. The batch is typically sampled at random from the entire dataset, so these statistics are intended to estimate their population-level counterparts [Ioffe, 2017]. At inference time, BN uses frozen statistics aggregated throughout the training process in place of batch statistics to standardize each test data point, one at a time.

We observed that prior preprocessing methods for correcting batch effects often standardardize data using an estimate of batch effects derived from averaging all samples in an experimental batch. For example, illumination correction for microscopy images divides images by the average intensity over all images from that plate [Singh et al., 2014]. Inspired by these methods, we hypothesized that calculating BN statistics specific to each experimental batch would similarly estimate batch effects, allowing these to be standardized out by the BN layers. The simplest way to achieve this is to change the data sampling process during training and inference. By ensuring that an input batch to the network is sampled from one experimental batch exclusively, we ensure that the calculated BN statistics reflects the means and variances of that experimental batch (rather than a global estimate of the entire population).

In practice, there are constraints and nuances in how this sampling is achieved. First, in principle, the batch size is important. With larger batch sizes, the BN statistics are a more precise estimate of the experimental batch effects. However, larger batch sizes may not always be feasible due to memory limitations. We solve this by using a smaller batch size during training, but not during inference. This works because backpropagating model gradients requires more memory during training, but no gradient tracking is needed during inference. Second, previous works have used different samples to estimate batch effects: some works use all data points [Singh et al., 2014], but other works use just control samples [Ando et al., 2017]. We argue that this decision depends on the availability of controls, but show that BEN can be effective both ways depending on the dataset. Pseudocode for BEN can be found in Appendix A.

## 3 Related Work

### Batch Correction Techniques

Normalization methods that correct for variations between different experimental batches have long been used within different areas of biological data analysis [Quackenbush, 2001, Redestig et al., 2009, Mecham et al., 2010, Leek et al., 2010, Piccinini et al., 2012]. Within microscopy, Singh et al. [2014] showed that using first-order statistics (i.e. means) to normalize images collected from a plate corrects for illumination. Later works [Ando et al., 2017] showed that incorporating second-order statistics (i.e. means and variances) into normalization led to effective correction of batch effects. These works have used these normalization methods during either data pre-processing or post-processing [Ando et al., 2017, Caicedo et al., 2018, Perakis et al., 2021, Moshkov et al., 2022]. In contrast, our work proposes normalization while training the neural network, and at every layer of the model’s internal feature representation (rather than just inputs or outputs).

In contrast to correction methods, some previous methods do directly correct for batch effects within training of a deep learning model. Qian et al. [2020] propose a generative adversarial network that equalizes images before training another network for morphological profiling. Wang et al. [2022] proposed correcting singlecell RNA-seq data using deep auto-encoders. Compared to these methods, which require training a separate network to tackle the batch effects problem prior to data analysis, BEN is simpler to implement and is not specific to any model set-up.

### Domain Adaptation using Batch Normalization

Adapting BN layers to improve generalization for neural networks has been studied in prior machine learning literature, most notably in domain adaptation [Li et al., 2017, Chang et al., 2019, Nado et al., 2020, Schneider et al., 2020, Zhang et al., 2021]. Our method resembles methods previously proposed by Chang et al. [2019] and Zhang et al. [2021], which use BN to normalize domains (similar to “experimental batches” in our setup). Chang et al. [2019] learn different BN parameters for each domain. In contrast, our method uses the same BN parameters for each domain, while only computing different BN statistics. As a result, our method can be applied to new experimental batches without re-training, and is more suited to high-throughput biology where new data is continuously generated. Zhang et al. [2021] introduce ARM-BN, a method similar to ours in terms of using domain-wise training and domain-wise inference. However, their empirical performance is worse than ours (Table 1). We identify two critical factors in our ablation studies: the use of a large batch size during inference, and correctly defining an “experimental batch”. Finally, most of these prior works focus on adapting BN for improving supervised classification models; our paper extends beyond supervised learning, showing that BEN enhances other machine learning settings routinely employed in microscopy, such as self-supervised learning and transfer learning [Ando et al., 2017, Hua et al., 2021, Perakis et al., 2021].

**Table 1:** Test set results on the RxRx1-Wilds supervised classification benchmark.

|  | Method | Top-1 Acc | Top-5 Acc |
| --- | --- | --- | --- |
| Our implementations | Vanilla | 31.1% | 50.5% |
|  | Vanilla + BEN | **40.2%** | **60.1%** |
| Wilds LeaderBoard | 1. IID Representation Learning [Wu et al., 2021] | 39.4% | N/A |
|  | 2. Vanilla + CutMix [Wu et al., 2021] | 38.4% | N/A |
|  | 3. LISA [Yao et al., 2022] | 31.9% | N/A |
|  | 4. ARM-BN [Zhang et al., 2021] | 31.2% | N/A |
|  | 5. Vanilla [Koh et al., 2021] | 29.9% | N/A |

## 4 Results

### 4.1 BEN results in state-of-the-art generalization for supervised learning

#### RxRx1-Wilds Dataset

First, we sought to understand if BEN could improve supervised learning on the RxRx1-Wilds dataset [Taylor et al., 2019, Koh et al., 2021], a large-scale fluorescence microscopy dataset used to understand batch effects. RxRx1-Wilds contains 125,514 images with size 256 × 256 and three fluorescent channels (nuclei, endoplasmic reticulum, actin) collected across 51 experiments. Each experiment involves cells from one of four cell lines (HUVEC, RPE, HepG2, or U2OS) exposed to the 1,139 different siRNA treatments. Each experiment uses four plates, each with 308 usable wells. Two images are taken from each well. The dataset proposes a classification task in which models are tasked with predicting what siRNA the cells were treated with, given an image. Importantly, the dataset splits are determined by experiment ID; of the 51 experiments, 33 are used for training, 4 are used for validation, and 14 are used for testing. Thus, for a model to perform well on the classification task, it must be able to overcome batch effects and generalize to experiments unseen during training.

#### Classification Model

Since our primary goal is to demonstrate the added benefit of BEN, we adopt the default modeling setup of the RxRx1-Wilds benchmark [Koh et al., 2021], which has optimally tuned the training hyperparameters of a ResNet50 classifier on this dataset. The classifier is trained with cross entropy loss for 90 epochs using the Adam optimizer [Kingma and Ba, 2015] (10^−3^ learning rate, 10^−5^ weight decay). The learning rate follows a linear warmup schedule for the first 10 epochs followed by a cosine annealing schedule for the final 80 epochs. The batch size during training is set to 75 images.

First, we trained a vanilla ResNet50 classifier. We report top-1 and top-5 accuracy in Table 1. Next, we applied BEN to the this model, as described in Section 2, using plate metadata provided by RxRx1-Wilds.

We observe that the use of BEN leads to increases of nearly 10% in both top-1 and top-5 accuracy on the RxRx1-Wilds supervised learning task. This result outperforms the state-of-the-art methods for this benchmark^2^. Unlike the top three existing methods (IID Representation Learning, Vanilla + CutMix, and LISA), our method does not utilize advanced data augmentation strategies (e.g. MixUp [Zhang et al., 2018], CutMix [Yun et al., 2019]) or multiple neural architectures (e.g. Restyle Encoder [Alaluf et al., 2021], StyleGAN Decoder [Karras et al., 2020]); all that BEN requires is a change in how we batch together data samples for training and evaluation.

To assess the impact of BEN qualitatively, we produced t-SNE plots colored by experimental batch (Appendix B.1.1). We observed that while the features learned by a baseline method cluster by experimental batch, this clustering is not evident in features learned with BEN.

### 4.2 BEN generalizes to improve self-supervised learning performance

Next, we sought to assess if BEN also improves unsupervised representation learning. Unsupervised strategies are popular in microscopy because they do not assume prior knowledge of biology [Caicedo et al., 2017, Lu et al., 2019b]. Despite the absence of labels, the goal is to learn representations that group together similar biological phenotypes, while distinguishing them from distinct phenotypes. Among the most effective strategies are self-supervised learning strategies, which propose unsupervised proxy tasks that teach skills that transfer to downstream analyses. Because there are no labels to guide the extraction of biological signal, unsupervised representation learning methods are also (if not more) susceptible to batch effects.

#### SimCLR

Previous work in applying self-supervised learning suggests that *simple contrastive learning* (SimCLR) – a method that trains models to maximize similarity between encoded representations of two augmented “views” of the same image [Chen et al., 2020] – is effective for morphological profiling, achieving strong performance in predicting the mechanism-of-action of drugs [Perakis et al., 2021]. Thus, we start with SimCLR as our baseline method.

To implement SimCLR on the RxRx1-Wilds dataset, we first segmented the images into single cell crops, in line with previous work [Perakis et al., 2021]. Each RxRx1-Wilds image was segmented into 48 × 48 cell crops. (See Appendix C.1 for segmentation details). Then, we trained a SimCLR model in a fully unsupervised fashion to obtain representations for each cell crop. Following Caicedo et al. [2017], we averaged the representation of every single cell in an image to form a representation of each image. To assess if SimCLR extracts information predictive of treatments, we fit a simple linear classifier on the image-level representations, following the same dataset splits used in the supervised evaluation (Section 4.1).

To generate views for SimCLR, cells were augmented using random rotations, random horizontal flips, random resized crops, and random Gaussian blurs. Though the original SimCLR algorithm uses two views per image, we found that using more than two views per cell resulted in better performance. In our experiments, we employed *K* = 5 views. Details on how to extend the SimCLR loss to *K >* 2 views, along with other implementation details (i.e. model architecture, training process) can be found in Appendix C.3.

Table 2 presents the treatment classification ability of SimCLR representations with and without BEN. In the “vanilla” SimCLR setup, each batch of cells was sampled *randomly* during training, by sampling images at uniform and then taking a random cell crop per image. When applying BEN, each batch of cells was sampled *plate-wise*, by first choosing a plate and then taking one cell crop from every image from that plate. Since the RxRx1-Wilds dataset has approximately 300 images per plate, we also set the training batch size to 300 images for the “vanilla” setup for fair comparison. During inference, BEN unfreezes the BN layers and performs inference using all cells in a plate as the inference batch, while the “vanilla” setup uses the default BN implementation. In Table 2, we observe that simply applying BEN to SimCLR results in significant improvements in predicting siRNA treatments, nearly doubling the top-1 accuracy of the linear classifier.

**Table 2:** RxRx1-Wilds test set results for linear classification of aggregated cell-level representations.

| Representations | Learning Algorithm | Vanilla (without BEN) |  | Our Method (with BEN) |  |
| --- | --- | --- | --- | --- | --- |
|  |  | Top-1 Acc | Top-5 Acc | Top-1 Acc | Top-5 Acc |
| Supervised | Single-Cell Classifier | 19.5% | 35.0% | **27.9%** | **47.3%** |
| Unsupervised | Random Network | 0.0% | 0.0% | N/A | N/A |
| Unsupervised | ImagetNet Classifier | 3.4% | 8.7% | N/A | N/A |
| Unsupervised | SimCLR ($K = 5$) | 11.0% | 22.9% | 19.0% | 35.5% |
| Unsupervised | MinCLR ($K = 5$) | 13.6% | 27.0% | **25.7%** | **44.1%** |

#### Baselines

Our set-up for unsupervised learning differs from the previous section in that we focus on single cells, not full images. The problem may be more challenging at the single cell level for many reasons, including loss of cells during segmentation, incomplete penetrance of phenotypes (not all cells may respond evenly to siRNA treatment) and information bottlenecks in the average pooling of single cells. Thus, we produced baselines at the single-cell level to provide further context in Table 2. As unsupervised baselines, we evaluated representations (i.e. at the final layer before the classification head) extracted by a ResNet50 model with randomly initialized weights, and with ImageNet initialized weights. Since these are methods that initialize fixed weights, rather than learning weights, BEN is inapplicable to these methods. The relatively low classification accuracies for these baselines in Table 2 confirms the difficulty of our unsupervised learning task; for several other tasks within the microscopy literature, these two baselines typically perform much better [Hua et al., 2021].

We also implement a supervised single-cell baseline. We used a ResNet50 classifier to classify cell crops according to their image-level siRNA treatment label during training and then discarded the final layer during inference to produce representations for each cell crop. We used the same training scheme (e.g. number of epochs, optimizer, batch size) as we did for SimCLR. We then fit a linear classifier to predict treatments at the image level, following the same procedure as the SimCLR representations. In Table 2, we observe that consistent with the more challenging nature of single-cell learning, supervised learning on the single-cell level underperforms full image classification. However, using BEN still results in large performance gains for the supervised single-cell baseline.

#### MinCLR

Since we observed a significant performance gap between off-the-shelf self-supervised methods (SimCLR) and supervised learning at the single-cell level, we decided to develop a new self-supervised method leveraging domain knowledge in fluorescence microscopy data, *multiple instance contrastive learning* (MinCLR). Since different cells from the same image were exposed to the same treatment, they can be considered “views” of each other (previous works exploited this assumption for multiple instance learning [Kraus et al., 2016, Lu et al., 2019a]). Instead of constructing views of a cell by randomly augmenting the same cell (as in SimCLR), MinCLR uses *different* cells from the same image as views of that cell. MinCLR encourages cells with the same treatment to have similar representations, making each representation robust to single-cell variation. Further details on MinCLR can be found in Appendix C.4. In our implementations, MinCLR uses the same data augmentations, modeling details, and training scheme as SimCLR. Like SimCLR, MinCLR also uses *K* = 5 views.

The performance of MinCLR representations with and without BEN is presented in Table 2. Similar to the SimCLR results, the MinCLR results show that using BEN results in large improvements in treatment classification accuracy, leading to almost a doubling in top-1 accuracy. Furthermore, MinCLR outperforms SimCLR and is closer to supervised performance. Scatter plots showing the effect of BEN on MinCLR representations can be found in Appendix B.1.3. We find that BEN aligns the representation spaces of different batches for MinCLR in a manner similar to what it does for SimCLR. However, unlike SimCLR + BEN, MinCLR + BEN appears to be qualitatively better at preserving certain biological features, such as cell line identities, in the representation space (Appendix B.2).

### 4.3 Ablation studies reveal factors important to BEN’s performance

#### 4.3.1 Batch-wise training and batch-wise inference are both crucial components

As described in Section 2, the two main components of BEN are batch-wise training and batch-wise inference (with unfreezing of BN layers). In Table 3, we present the impact of switching one or both of these components to standard BN practice. “Vanilla training” trains on batches randomly sampled from the entire dataset, and “vanilla inference” involves evaluating each test image using frozen BN statistics computed during training.

**Table 3:** Ablation study for linear prediction of MinCLR representations on the RxRx1-Wilds test set.

| Method | Top-1 Acc |  |
| --- | --- | --- |
| | $K = 2$ | $K = 5$ |
| MinCLR + (vanilla training, vanilla inference) | 12.5% | 13.6% |
| MinCLR + (vanilla training, batch-wise inference) | 16.1% | 16.9% |
| MinCLR + (batch-wise training, vanilla inference) | 10.6% | 15.3% |
| MinCLR + BEN (i.e. batch-wise training, batch-wise inference) | **18.7%** | **25.7%** |

The ablation is performed on MinCLR, the best unsupervised learning algorithm from Section 4.2. MinCLR has a hyperparameter *K*, which controls the number of views; we reported top-1 classification accuracy for both *K* = 2 and *K* = 5.

We observed that while either batch-wise training alone or batch-wise inference alone led to some performance improvements, their combination resulted in the best performance. These results support our hypothesis that both components synergize, with batch-wise training removing experimental batch effects during training, and batch-wise inference doing the same for data from new experimental batches during evaluation. When both are used, the model is trained *and* evaluated on corrected data, leading to the best performance.

#### 4.3.2 BEN requires correct specification of “batch” and large batch sizes during inference

Next, we found that (1) the definition of an experimental batch and (2) the use of the full batch during inference were critical to BEN’s performance. Following previous literature on plate effects [Caicedo et al., 2017], we chose to use a “plate” as the experimental batch throughout our results in Section 4. However, the RxRx1 dataset contains metadata for experiments, plates, wells, and imaging sites, and previous similar works exploiting BN for domain adaption consider a domain to be an experiment instead [Zhang et al., 2021]. Here, we find that using plates, rather experiments, leads to superior performance (Table 4).

**Table 4:** Ablation study results on the RxRx1-Wilds supervised classification benchmark.

| Method | Top-1 Acc |
| --- | --- |
| Vanilla + BEN (i.e. batching by plate, full batches during inference) | **40.2%** |
| Vanilla + (batching by plate, minibatches during inference) | 36.9% |
| Vanilla + (batching by experiment, full batches during inference) | 35.1% |
| Vanilla + (batching by experiment, minibatches during inference) | 33.7% |
| Vanilla (i.e. vanilla batching, vanilla inference) | 31.1% |

Second, consistent with our hypothesis that a larger sample size during inference would result in a better estimation of batch effects, we observed that using the entire experimental batch during inference (as opposed to using a batch size consistent with training) also led to improvements in performance. Table 4 presents our findings for supervised classification on the RxRx1-Wilds image-level dataset, following the set-up in Section 4.1. When using batches during inference, we set the batch size to 75 to be consistent with training.

### 4.4 BEN leads to effective transfer learning between different microscopy datasets

Finally, we explored the impact of BEN on transfer learning. In morphological profiling, one practice is to use a pre-trained model as an off-the-shelf feature extractor. Unlike supervised or unsupervised representation learning, transfer learning does not require training a model at all, making it a practical solution for labs with less compute resources [Ando et al., 2017, Pawlowski et al., 2016, Hua et al., 2021].

Following this setting, our aim is to use a model pretrained on one dataset as a feature extractor for another, distinct dataset (e.g. one collected by a different lab with different markers). To evaluate this, we train single-cell models on the RxRx1-Wilds dataset and evaluate on the BBBC021 dataset [Caie et al., 2010], a morphological profiling benchmark for drug discovery. We compare our models against methods that employ a similar transfer learning setup. While there have been several works that first pre-train a model on BBBC021 [Godinez et al., 2017, Caicedo et al., 2018, Perakis et al., 2021] prior to evaluation, these works train on the dataset of interest, and are therefore not the main focus of comparison.

We present results in Table 5. We transfer models trained using MinCLR (Section 4.2) on RxRx1-Wilds. We apply typical variation normalization (TVN) to our features, a standard post-processing step for this dataset first introduced by Ando et al. [2017]. Following prior practice [Ando et al., 2017], we report not-same-compound (NSC) and not-same-compound-batch (NSCB) accuracy, which measure correct assignment of mechanism of action using a nearest-neighbor classifier, controlling for compound and batch labels. Further details on modeling, implementation, the effect of TVN, and learned drug representations can be found in Appendix D. We compare results against our own baselines (random features and ImageNet features) and those from the literature. We have three main observations: (1) Applying BEN boosts the performance of transfer learning. (2) For this dataset, better performance is achieved when *control samples* are used to compute normalization statistics within BN layers (as opposed to all samples). This is consistent with other works for this dataset that also only use control samples for post-processing normalization [Ando et al., 2017, Caicedo et al., 2018]. (3) Prior to this work, there were significant gaps in transfer learning performance between models trained on microscopy images (i.e. CytoImageNet [Hua et al., 2021]) and models trained on natural images. We show that MinCLR + BEN trained on RxRx1-Wilds closes this gap for NSC accuracy.

**Table 5:** BBBC021 1-nearest-neighbor mechanism-of-action classification results using transfer learning.

|  | Dataset | Architecture | Learning Algorithm | NSC | NSCB |
| --- | --- | --- | --- | --- | --- |
| Ours | N/A | ResNet50 | Random Network | 77% | 61% |
|  | ImageNet | ResNet50 | Classifier | 85% | 72% |
|  | RxRx1-Wilds | ResNet50 | MinCLR (no BEN) | 84% | 78% |
|  | RxRx1-Wilds | ResNet50 | MinCLR + BEN (BN w/ all samples) | 92% | 75% |
|  | RxRx1-Wilds | ResNet50 | MinCLR + BEN (BN w/ controls) | **96%** | 88% |
| Literature | ImageNet | Inception | Classifier [Pawlowski et al., 2016] | 91% | N/A |
|  | Product Images | AlexNet | Metric Learning [Ando et al., 2017] | **96%** | **95%** |
|  | N/A | N/A | Classical Features [Caicedo et al., 2018] | 90% | 85% |
|  | CytoImageNet | EfficientNet | Classifier [Hua et al., 2021] | 86% | N/A |

## 5 Conclusion

In this work, we introduced *batch effects normalization* (BEN), a simple method for correcting batch effects that can be applied to anyx neural network with batch normalization layers. We demonstrated that BEN led to significant improvements in performance across supervised learning, self-supervised learning, and transfer learning for microscopy images, resulting in state-of-the-art performance on the RxRx1-Wilds dataset.

We believe that there are several opportunities for future work. First, future work should probe how the definition of an experimental batch interacts in greater detail. Although we defined an experimental batch as a plate (and showed the superiority of this strategy over definitions that group multiple plates together), wells in one quadrant of a plate may experience different effects than wells in another quadrant of the same plate [Caicedo et al., 2017]. Second, though the batch size during inference cannot be too small, it is unclear if the full experimental batch is always needed, or if there is a “break point” in which a large enough batch size leads to the same performance. Third, in all of our experiments with BEN, we constructed training batches by sampling exclusively from one experimental batch. This could be limiting for datasets with small experimental batches and it would be interesting to see if BEN could be generalized to simultaneously train on samples from different experimental batches. We leave these as open questions for future investigation.

## Acknowledgements

We thank Juan C. Caicedo, Anne E. Carpenter, Shantanu Singh, Kianoush Falahkheirkhah, Wenjun Wu, and the BioML team at Microsoft Research for their very helpful feedback.

## A Appendix: Pseudocode for Typical BN and BEN

Assume a training dataset 𝒳_train_ is collected in experimental batches 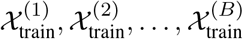. The test dataset consists of data from new batches 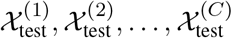 that were unseen during training. In Algorithm 1, we review the typical deep learning pipeline that is used with batch normalization (BN) layers. In Algorithm 2, we summarize how BEN changes this pipeline with the goal of correcting for batch effects. In our notation for this section, the operations +,−, *, /, √ are all applied element-wise on vectors, where appropriate. As a final note, BEN optionally allows for computing BN statistics using control samples, following other methods that use controls for normalizing data [Ando et al., 2017]. This is applicable in datasets where each experimental batch 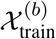(resp. 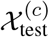contains a subset of labeled controls 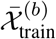(resp. 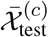.

### Algorithm 1

Typical Deep Learning with Batch Normalization (parameters: ***β, γ***; hyperparameters: *δ, ϵ*)

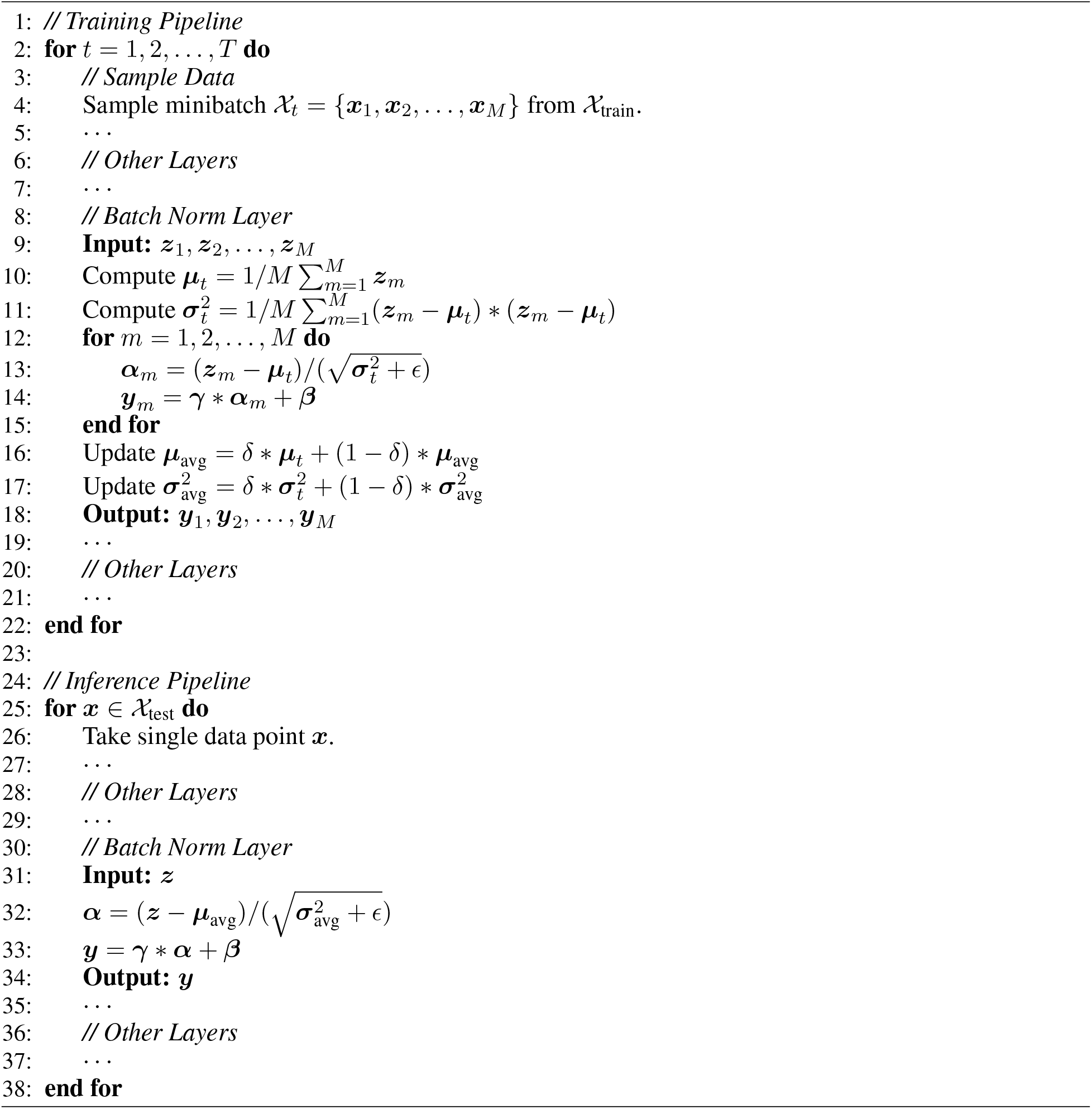

### Algorithm 2

Batch Effects Normalization

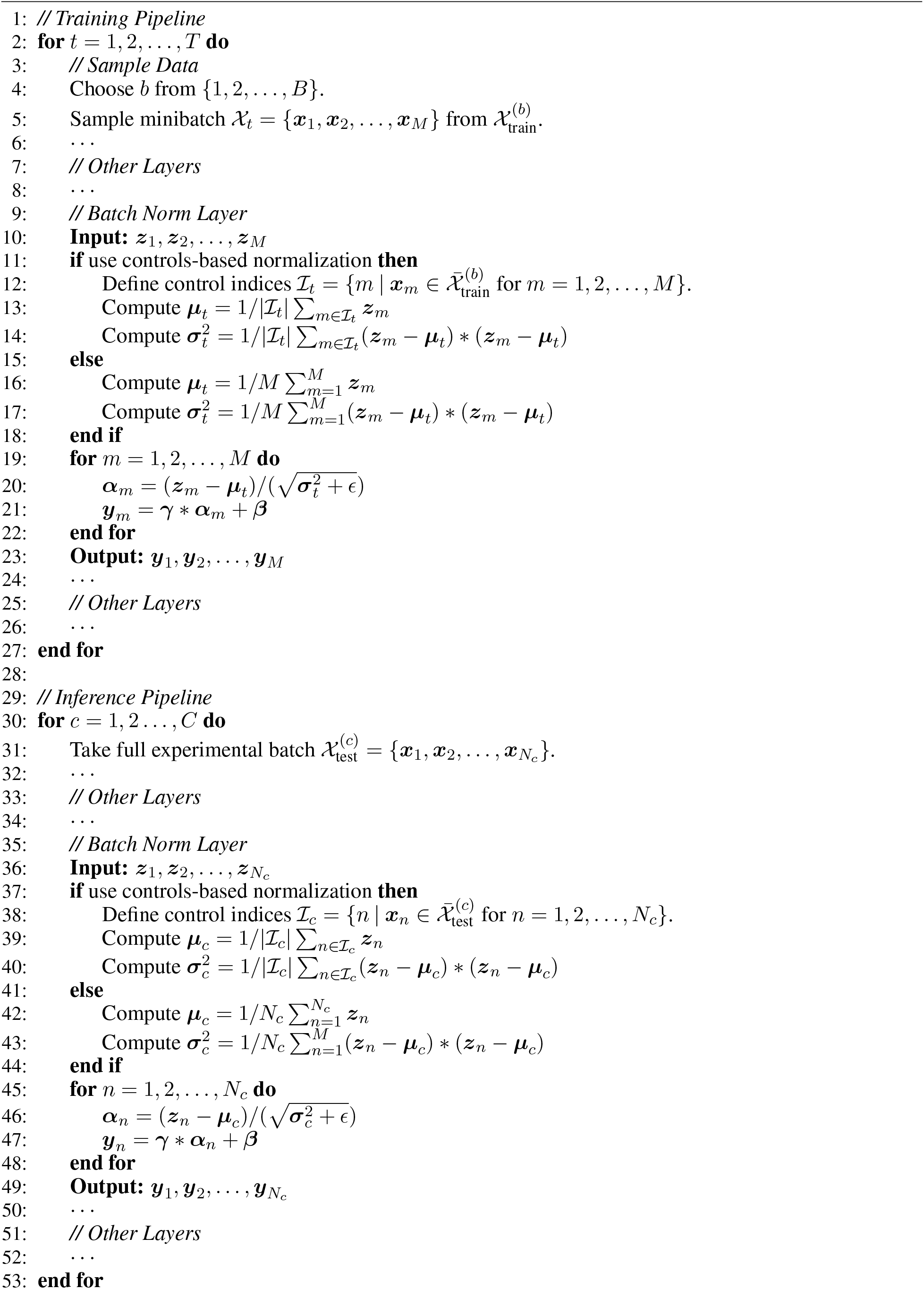

## B Appendix: Representation Space Figures

In these figures, we qualitatively analyze image-level vector representations learned by different models for microscopy images from the RxRx1-Wilds test set [Taylor et al., 2019, Koh et al., 2021]. The original *D* = 2048-dimensional vectors are projected down to two dimensions using t-SNE [Van der Maaten and Hinton, 2008] and colored by their RxRx1-Wilds experiment ID. Each experiment consists of images of cells from a particular cell line exposed to the same 1,139 siRNA treatments. In each figure in this section, we compare the representations learned by a standard deep neural network and those learned by the same network when BEN is applied.

### B.1 Repeated HUVEC Experiments

In this subsection, we visualize the representations of images from four repeated HUVEC experiments. Since the cells from the different experiments are from the same cell line (and the set of treatments is the same), we expect a network that is robust to batch effects to not have significant differences in distribution among its image representations from different experiments. Across deep learning models trained using various supervised and self-supervised algorithms, we consistently find that models from the vanilla approach separate in the feature space according to experimental batch, whereas those that have applied BEN do not.

#### B.1.1 Supervised Image Representations

These representations are obtained from the image-level ResNet50 supervised classifier of Section 4.1. To obtain the representations, we remove the final classification layer of the ResNet and visualize the features in the penultimate layer for each image.

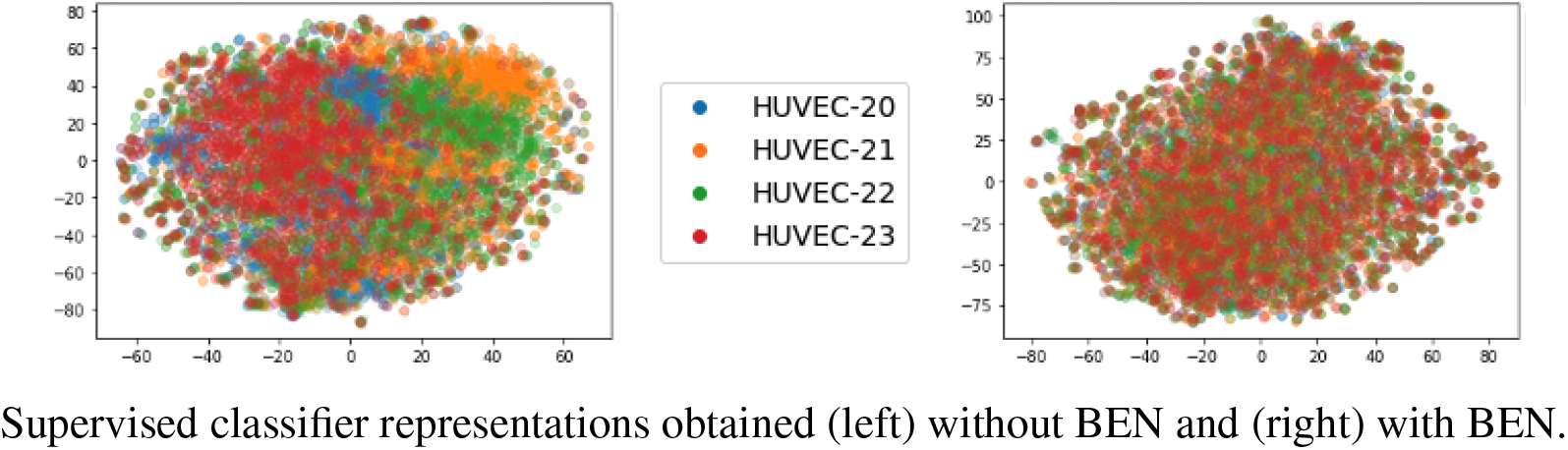

#### B.1.2 Pooled SimCLR Single-Cell Representations (Figure 1)

Figure 1 (reproduced below for convenience) visualizes image representations that were obtained using the SimCLR algorithm [Chen et al., 2020]. As described in Section 4.2, SimCLR is trained on single-cell crops. To obtain image-level representations, we take an average pooling of representations for all cropped cells within an RxRx1-Wilds image. While training SimCLR, we use *K* = 5 views per cell.

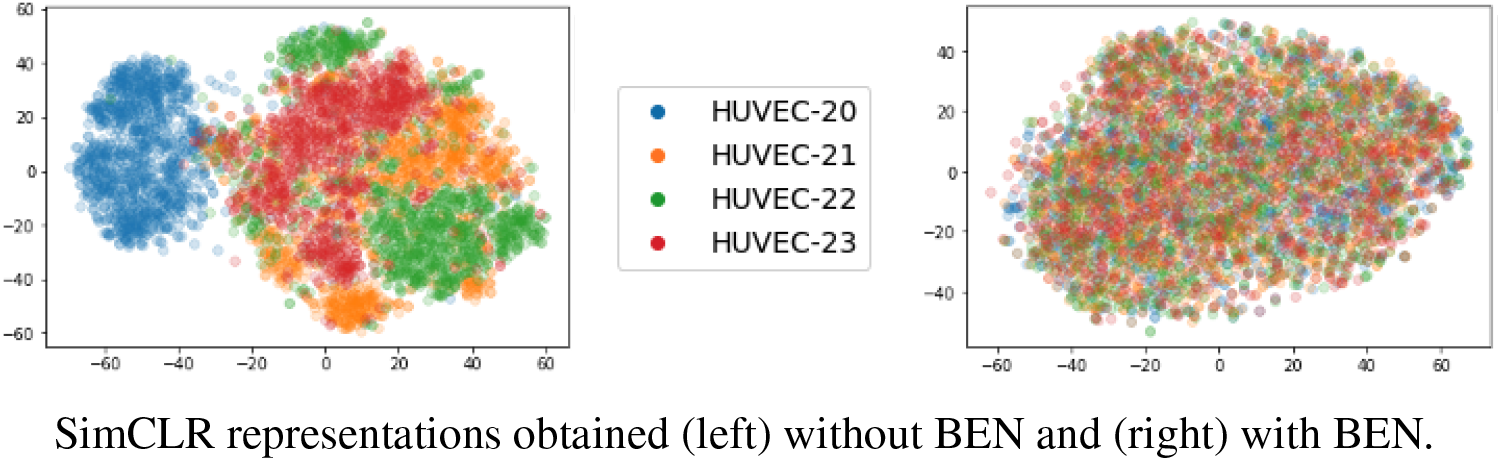

#### B.1.3 Pooled MinCLR Single-Cell Representations

Here, cell-level representations are first trained using MinCLR, as described in Section 4.2, and then aggregated in the same way as SimCLR representations. Like SimCLR, we use *K* = 5 views per cell.

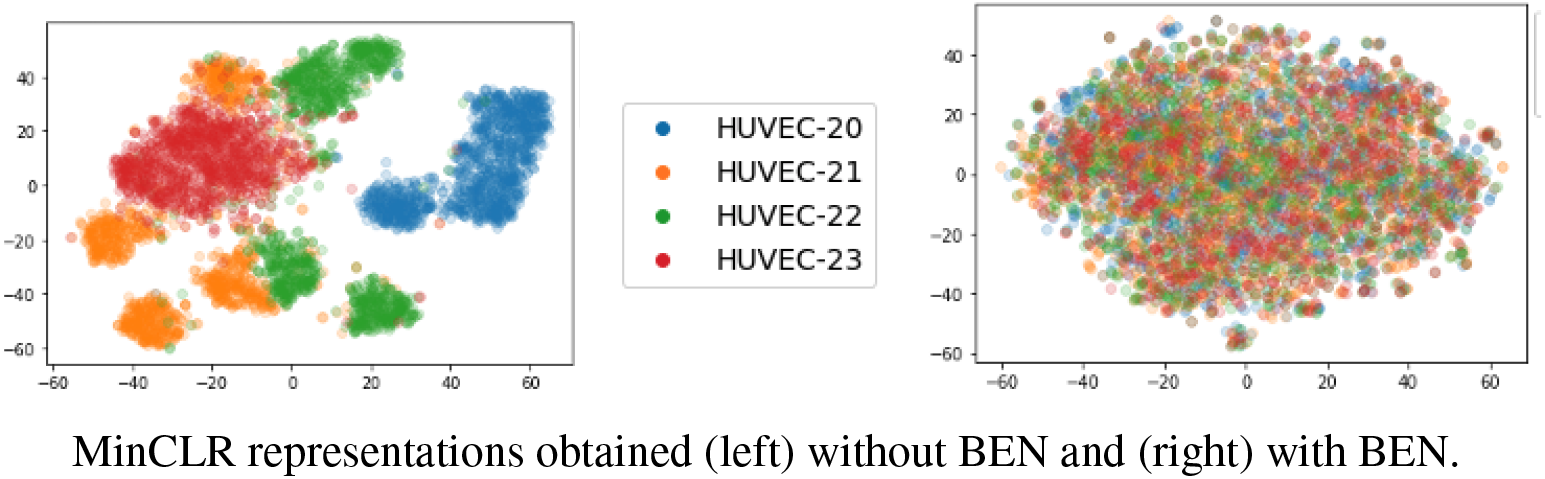

#### B.1.4 Pooled Supervised Single-Cell Representations

Here, we first trained a supervised single-cell treatment classifier, as described in Section 4.2. We then removed the final classification layer of the model and used the penultimate layer to obtain representations for each single-cell crop. Then, these representations are aggregated to form image-level representations in the same way as SimCLR and MinCLR.

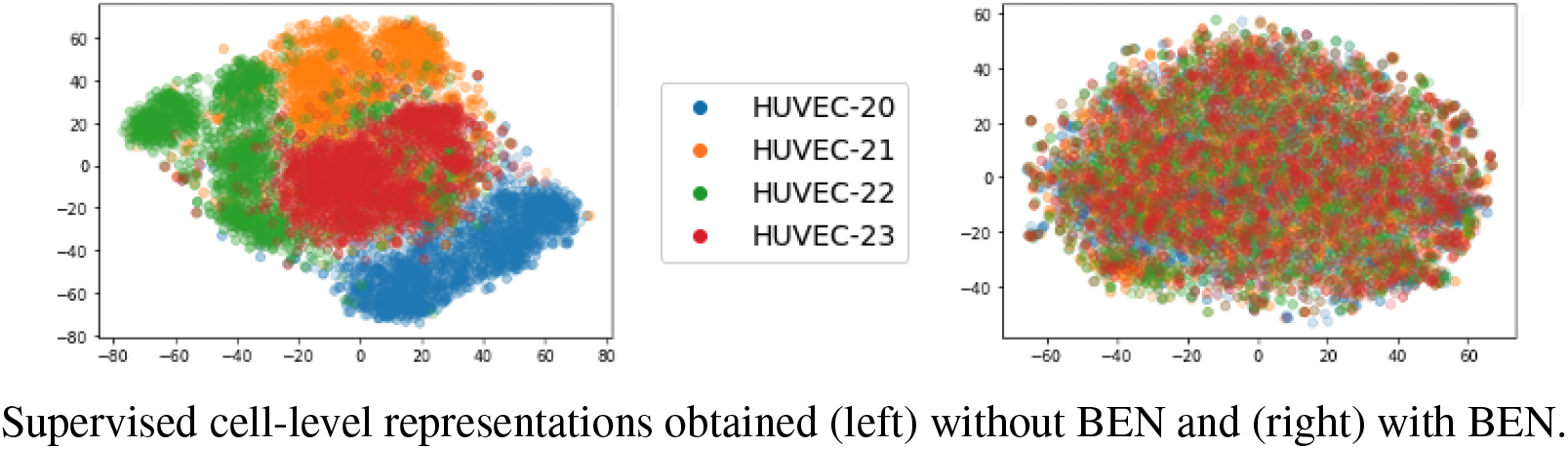

### B.2 Different Cell Lines

In this subsection, we visualize image representations from experiments involving *different* cell lines. Specifically, we look at 2 HUVEC experiments, 2 HepG2 experiments, 2 RPE experiments, and 1 RPE experiment (since there is only a single RPE experiment in the RxRx1-Wilds test set). We examine representations from the SimCLR and MinCLR self-supervised learning algorithms (Section 4.2). From the figures below, we have two main observations: (1) For both SimCLR and MinCLR, using BEN removes differences between experiments of the same cell line. (2) MinCLR + BEN preserves distributional differences *between cell lines* better than SimCLR + BEN (which appears to merge together RPE and HUVEC representations).

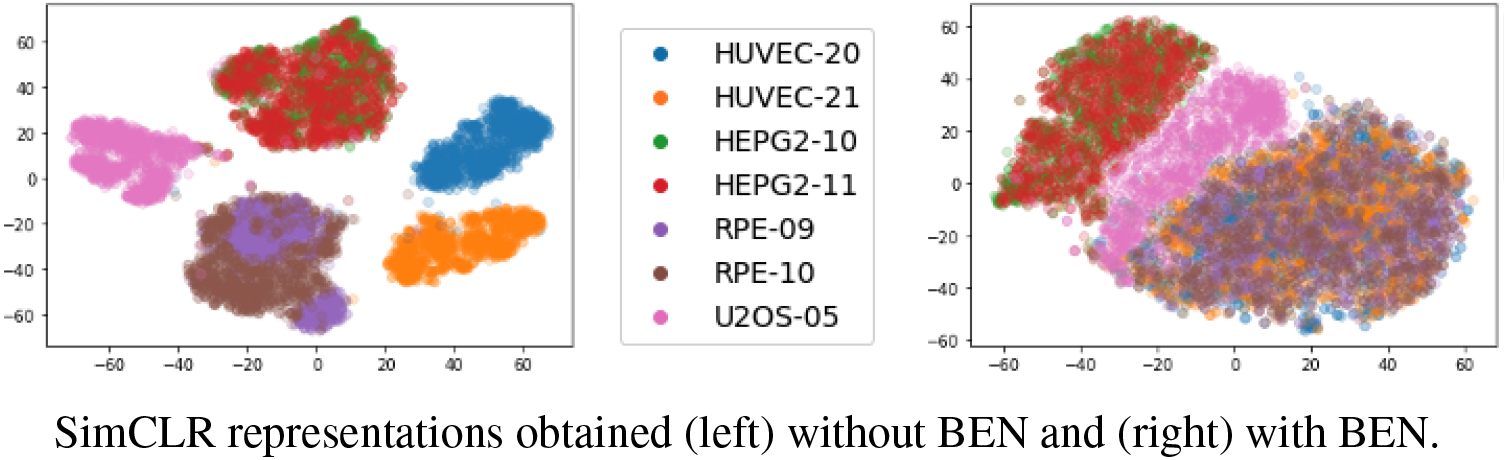

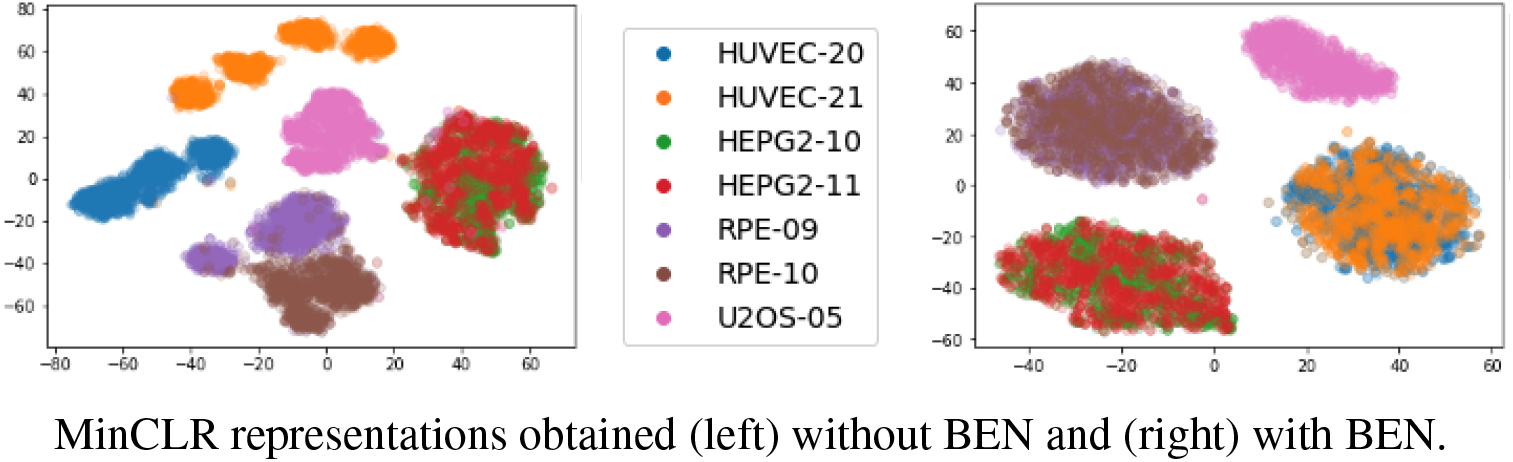

## C Appendix: Additional Details for Self-Supervised Learning

### C.1 Cell-Level RxRx1-Wilds

To create a dataset of single cell images from RxRx1-Wilds [Taylor et al., 2019, Koh et al., 2021], we segmented the nuclei channel of each 256 × 256 image using a Mask R-CNN model [He et al., 2017]. The segmentation model was pretrained on the 2018 Data Science Bowl [Caicedo et al., 2019], a dataset of over 37,000 manually annotated nuclei^3^. Given each segmented mask, we made a 48 × 48 pixel crop around the mask’s center of mass. This crop was discarded if the mask had an area that was less than 10% of the crop region, or if the cropped region exceeded the image boundaries. All crops that were not discarded were applied to the three channels of the original image to generate a single cell image. Note that due to crowding of cells within the original image, some of our single cell images may actually contain more than one cell, even though they are always centered around one cell. In the figure below, we present some statistics on the number of cells segmented per image for the RxRx1-Wilds training set, broken down by different cell lines.

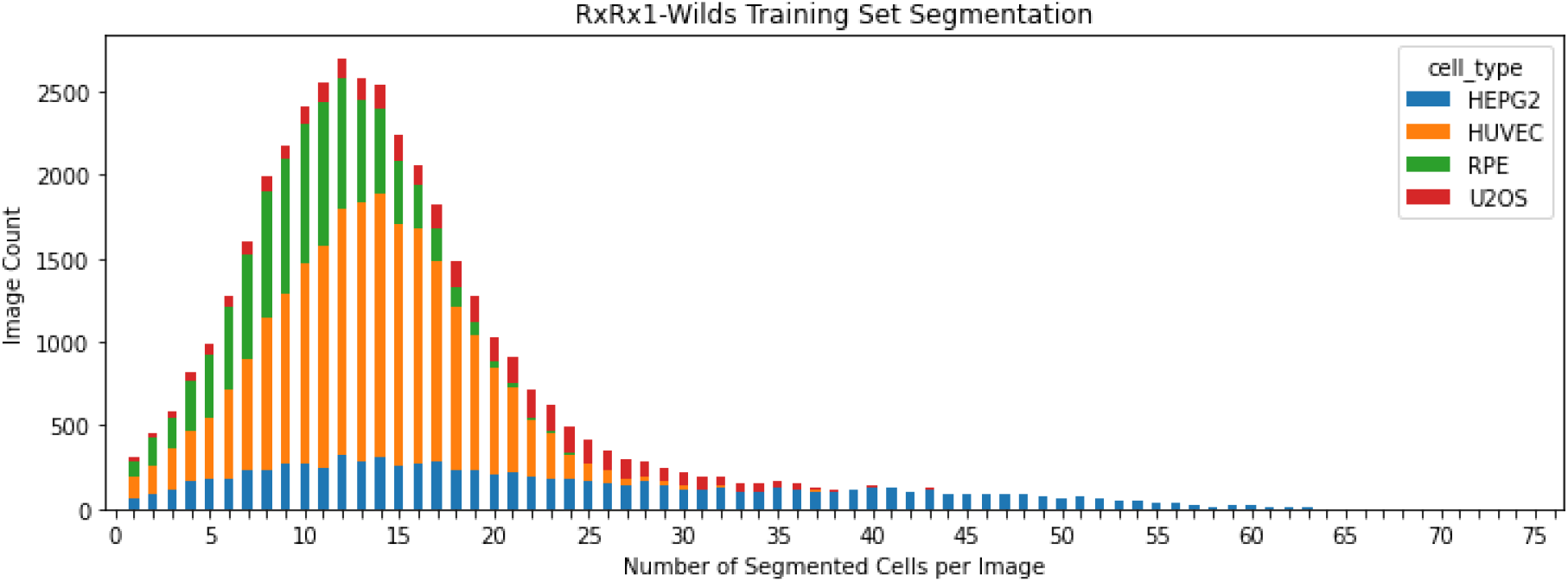

### C.2 Review of Standard SimCLR

To train the standard SimCLR algorithm [Chen et al., 2020] on the RxRx1-Cells dataset, we form each minibatch at training step *t* by first sampling a set of images 𝒳_t_ from RxRx1-Wilds. Then, for each ***x*** ∈ 𝒳_t_, we randomly sample a cell ***c*** ∈ 𝒞^***x***^, where 𝒞^***x***^ denotes the set of all segmented cells in image ***x***. Using data augmentation, we create two views for each ***c***. We assemble these views into a cell-level minibatch 𝒞_t_ of size |𝒞_t_| = 2 · |𝒳_t_|. We pass each view 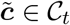 through the encoder *f* to generate a representation 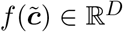. The representation is further passed through a nonlinear transformation *g* to yield a projection 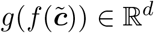. For each 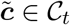, the SimCLR loss function trains its projection to attract the projection of its “positive anchor” 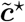 (i.e. the other view that originated from the same cell ***c***), and repel the projections of all other views in 𝒞_t_:

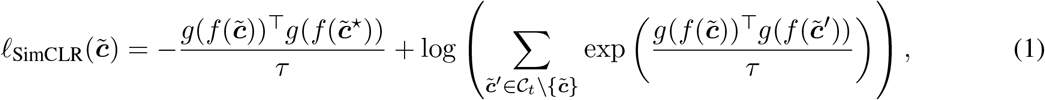

where *τ* ∈ ℝ is an adjustable temperature parameter. After training, the nonlinear projector *g* is discarded and *f* is used to generate representations *f* (**𝒸**) for each cell **𝒸**.

### C.3 Our Implementation of SimCLR

In the original SimCLR algorithm [Chen et al., 2020], two augmented views are used per image. However, we found that using *K >* 2 views typically resulted in better performance. Additional views can be generated for SimCLR by simply applying the random data augmentations more times. In these cases, the cell-level minibatch 𝒞 _t_ has size | 𝒞_t_ | = *K* · | 𝒳 _t_ |. Since there are now multiple (i.e. *K* − 1) positive anchors for each view 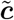, we need a more general loss function than Eq. (1). Let 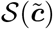 denote the set of positive anchors for 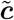. Drawing inspiration from Khosla et al. [2020], we propose the following *K*-view SimCLR loss:

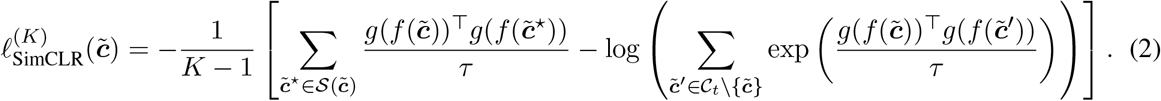

In Section 4.2, we used the following settings when implementing SimCLR: We set the SimCLR temperature parameter to 0.05. Following Chen et al. [2020], we use a ResNet50 with the final classification layer removed and replaced by a two-layer nonlinear projector (that is discarded after training) mapping *D* = 2048 dimensions to *d* = 128. The model was initialized using pretrained ImageNet weights and train for 500 epochs on the RxRx1-Wilds single cell dataset. We used the Adam optimizer with a learning rate of 10^−3^, linear warmup for the first 50 epochs, cosine annealing for the final 450 epochs, and no weight decay. To fit the linear classifier, we used minor data augmentations (i.e. random rotations and random horizontal flips), as well as the Adam optimizer with learning rate 10^−1^, no weight decay, and cosine annealing for 30 epochs.

### C.4 Mathematical Details for MinCLR

The only difference between MinCLR and SimCLR is how to define the views of each image. Whereas SimCLR augments the same cell using multiple augmentations, MinCLR augments multiple different cells from the same image.

Each minibatch at training step *t* for MinCLR is formed as follows: Like SimCLR, we first sample a set of images 𝒳_*t*_. Then, for each ***x*** _t_, we randomly sample *K* cells **𝒸**_1_, **𝒸**_2_, …, **𝒸**_*K*_ ∈ 𝒞 ^***x***^. The cells are sampled with replacement, because it may be the case that | 𝒞^***x***^ | *< K* for a particular ***x***. Next, we apply the same data augmentation as in SimCLR to generate augmented cells 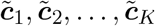. Let _t_ denote all the augmented cells within the MinCLR minibatch. It has size| 𝒞_t_; | = *K* · | 𝒳_t_ |. Each 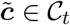 is then passed through the encoder *f* and the projector *g*. For each 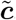, let 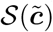 denote the set of *K* − 1 other augmented cells that originated from the same image ***x***. The *K*-view MinCLR loss function is:

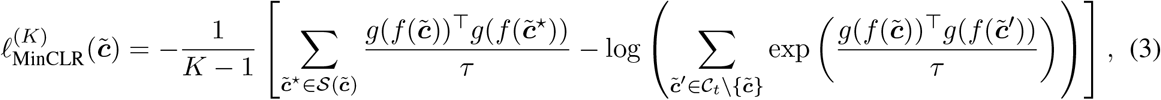

which is the same as Eq. (2), with the only difference being in how 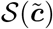 is defined.

## D Appendix: Additional Details for Transfer Learning

### D.1 Modeling and Evaluation Details

To train our cell-level feature extractor on RxRx1-Wilds, we employed the MinCLR algorithm from Section 4.2 with *K* = 5 views following the same implementation details as in that section. We chose this single cell model due to a mismatch in image size between the RxRx1-Wilds dataset and the BBBC021 dataset, making full image models more challenging to transfer. When training MinCLR + BEN (BN w/ control samples), we calculated BN statistics for each plate by only using the control samples, with control labels provided by the RxRx1-Wilds dataset. Similarly, for evaluation, we calculated the unfrozen BN statistics for each plate by using the control (i.e. DMSO) labels provided by BBBC021. Since RxRx1-Wilds and BBBC021 measure different fluorescent channels, we used the following channel mappings for the model: DNA → nucleus, tubulin → endoplasmic reticulum, actin → actin.

For BBBC021, we first downloaded all microscopy images using the pybbbc package, accessed at https://github.com/giacomodeodato/pybbbc. Following prior work [Perakis et al., 2021], we used the single-cell nuclei locations provided by Ljosa et al. [2013] to segment the images into 48 × 48 crops. Crops were discarded if they exceeded the image boundary or if the nucleus mask contained less than 250 pixels.

The goal of the BBBC021 dataset is to evaluate the whether 103 drugs (i.e. compound-concentration pairs) cluster by mechanism of action in a feature space representing images of MCF-7 breast cancer cells treated with these drugs. Following previous work [Caicedo et al., 2018, Perakis et al., 2021], we constructed representations for each of drugs as follows: For each cell crop, we obtained a vector representation using the pretrained cell-level feature extractor. Within an image, all cell-level representations were aggregated by taking an element-wise mean. Then, all image-level representations for a drug were aggregated by taking an element-wise median.

During evaluation, each drug representation was assigned the functional class (i.e. one of 12 mechanism-of-actions) of its closest neighbor by cosine distance. The mechanism-of-action accuracy is computed as the fraction of correct assignments based on the known functional classes. From the literature [Ando et al., 2017], there are two ways to evaluate this accuracy based how one defines the closest neighbor: For the not-same-compound (NSC) metric, the closest neighbor cannot come from drugs of the same compound. For the not-same-compound-batch (NSCB) metric, the closest neighbor cannot come from drugs of the same compound or of the same experimental batch. The NSCB metric is always calculated on a smaller subset (i.e. 92 of the 103 drugs) because certain classes of treatments were only collected in one particular batch.

### D.2 Additional Results

We report additional results for transfer learning, extending Table 5. Specifically, we show the effect of different post-processing steps during inference (i.e. no post-processing, whitening, and typical variation normalization [Ando et al., 2017]) and how they interact with BEN. Note that TVN involves a whitening step followed by correlation alignment (CORAL) [Sun et al., 2016]. The whitening step is used to equalize the scales of the features, while the CORAL step is used to correct for batch effects. See Ando et al. [2017] for more details.

We make the following observations from the table below: (1) When BEN is not used, both whitening and TVN lead to improvements over no post-processing. (2) When BEN is used, TVN provides no added improvements beyond whitening. (3) In addition, when TVN is used, BEN still provides added benefits over when BEN is not used. Taken together, these three observations appear to suggest that BEN subsumes the ability of CORAL to correct for batch effects, but not the other way around. We speculate that this is because the scope of BEN is much wider than that of CORAL – i.e. BEN affects the training process and every intermediate layer of a network during evaluation. On the other hand, CORAL is only applied as a post-processing step on the final representations.

**Table 6:** BBBC021 1-nearest-neighbor classification results using different types of post-processing.

| Dataset | Learning Algorithm | None |  | Whiten |  | TVN |  |
| --- | --- | --- | --- | --- | --- | --- | --- |
|  |  | NSC | NSCB | NSC | NSCB | NSC | NSCB |
| RxRx1-Wilds | MinCLR (no BEN) | 75% | 62% | 80% | 72% | 84% | 78% |
| RxRx1-Wilds | MinCLR + BEN (BN w/ all samples) | 86% | 69% | 92% | 75% | 92% | 75% |
| RxRx1-Wilds | MinCLR + BEN (BN w/ controls) | **95%** | **85%** | **96%** | **88%** | **96%** | **88%** |

### D.3 Drug Representations

In the figures below, we visualize the drug representations projected down to two dimensions using t-SNE for different cell-level feature extractors. The drugs are colored based on their known mechanism-of-action. Consistent with Table 5, we observe that the feature space learned by pretraining on RxRx1-Wilds with BEN places drugs in the same class closer together than pretraining on RxRx1-Wilds without BEN and pre-training on ImageNet.

**Figure 2:**
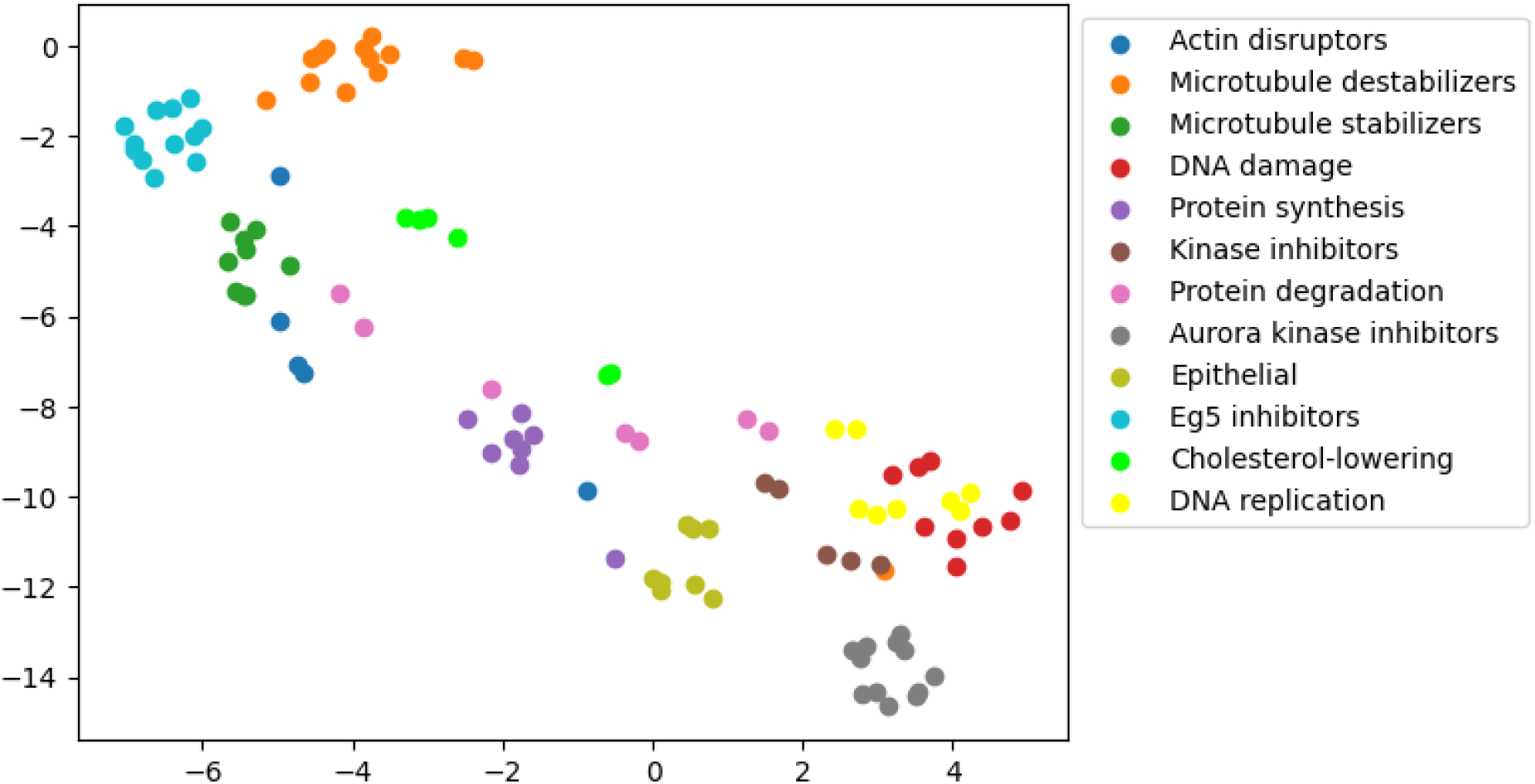
Representation space of drugs learned by pretraining on ImageNet.

**Figure 3:**
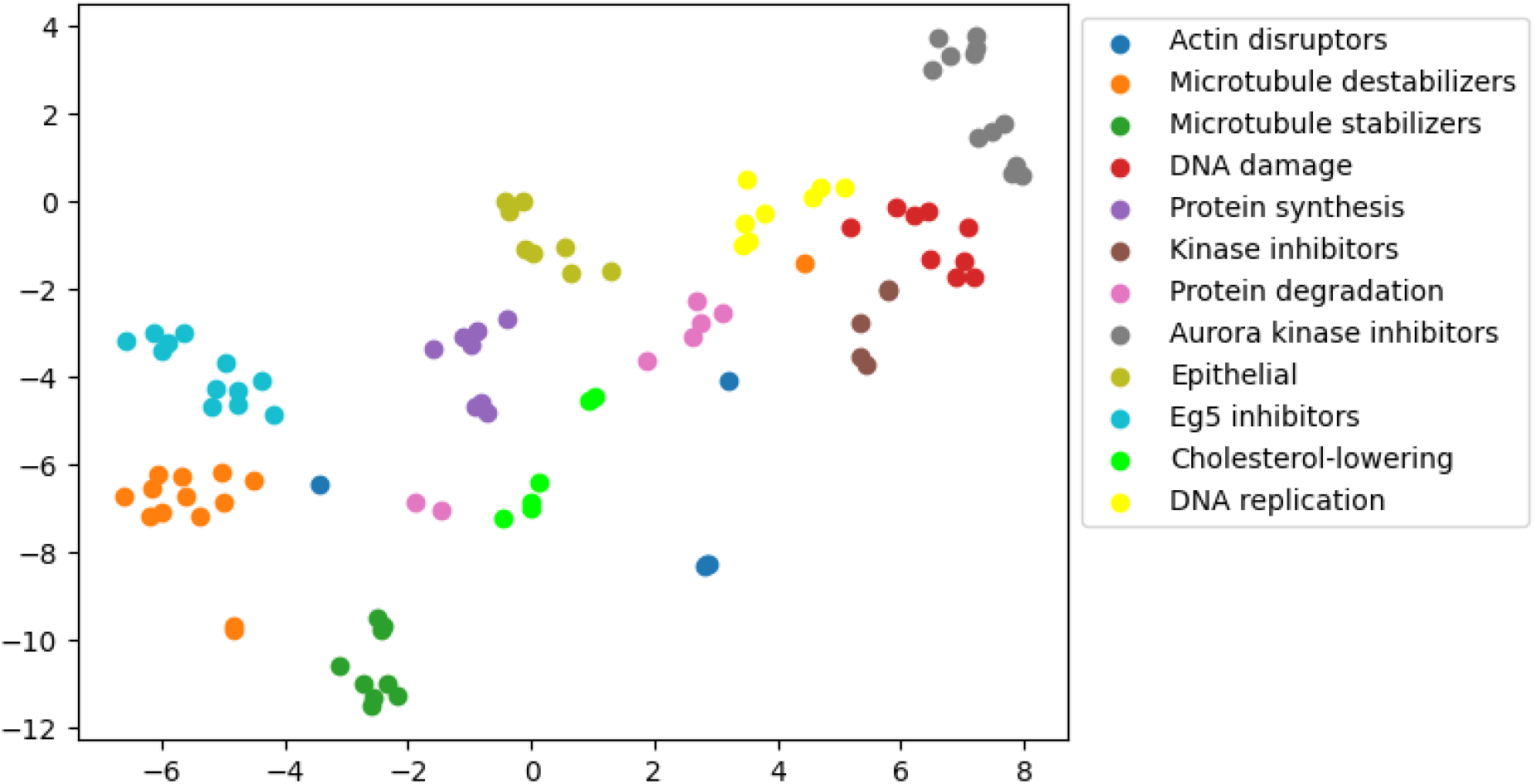
Representation space of drugs learned by pretraining on RxRx1 without BEN.

**Figure 4:**
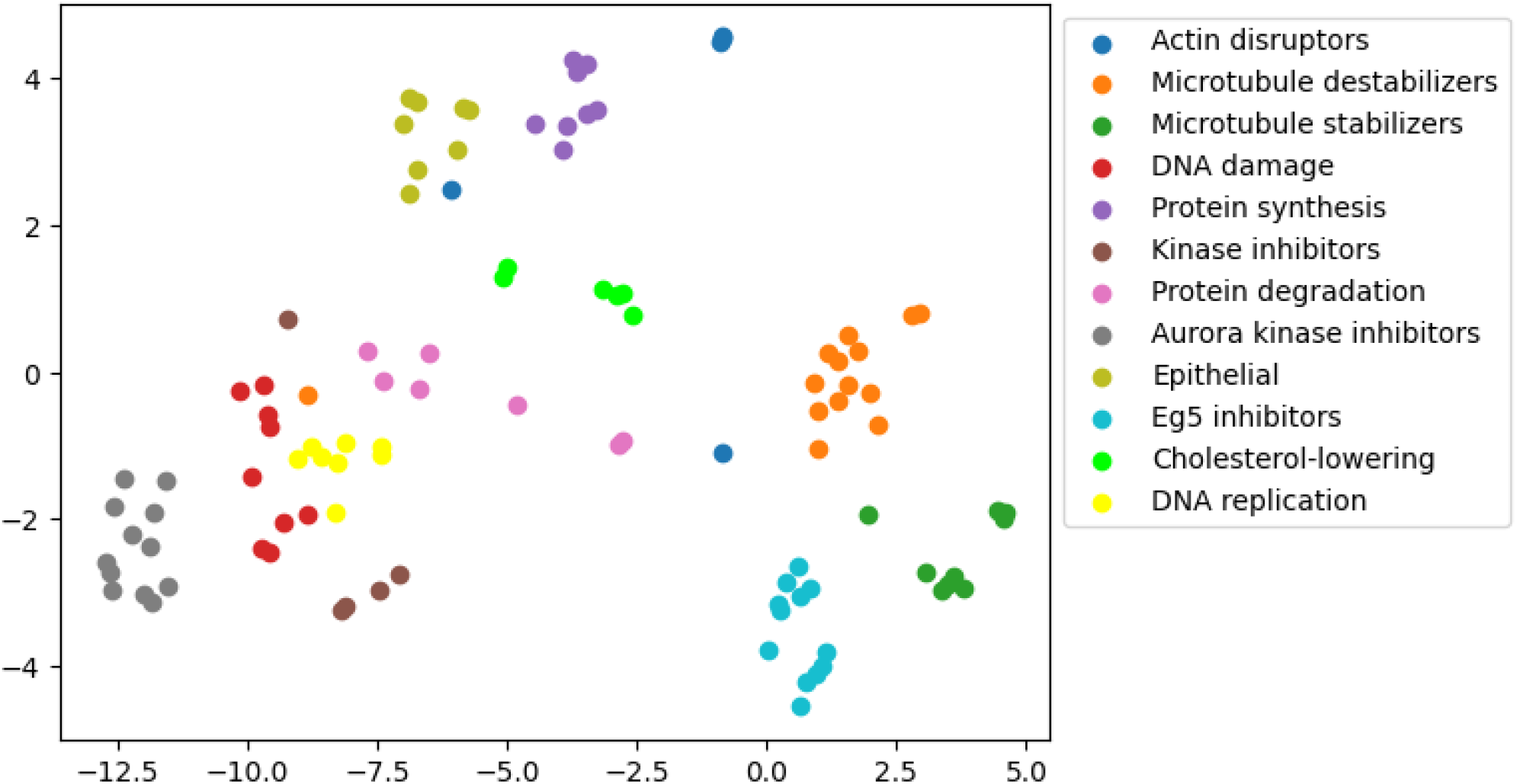
Representation space of drugs learned by pretraining on RxRx1 with BEN.

## Footnotes

1 Our code can be accessed at https://github.com/microsoft/batch-effects-normalization/.

2 Wilds LeaderBoard numbers were sourced from https://wilds.stanford.edu/leaderboard/.

3 The pretrained model can be accessed at https://github.com/Lopezurrutia/DSB_2018.

